## Supplementary material for "Striated muscle-specific base editing enables correction of mutations causing dilated cardiomyopathy": Sup. Figures

**Supplementary Figures**

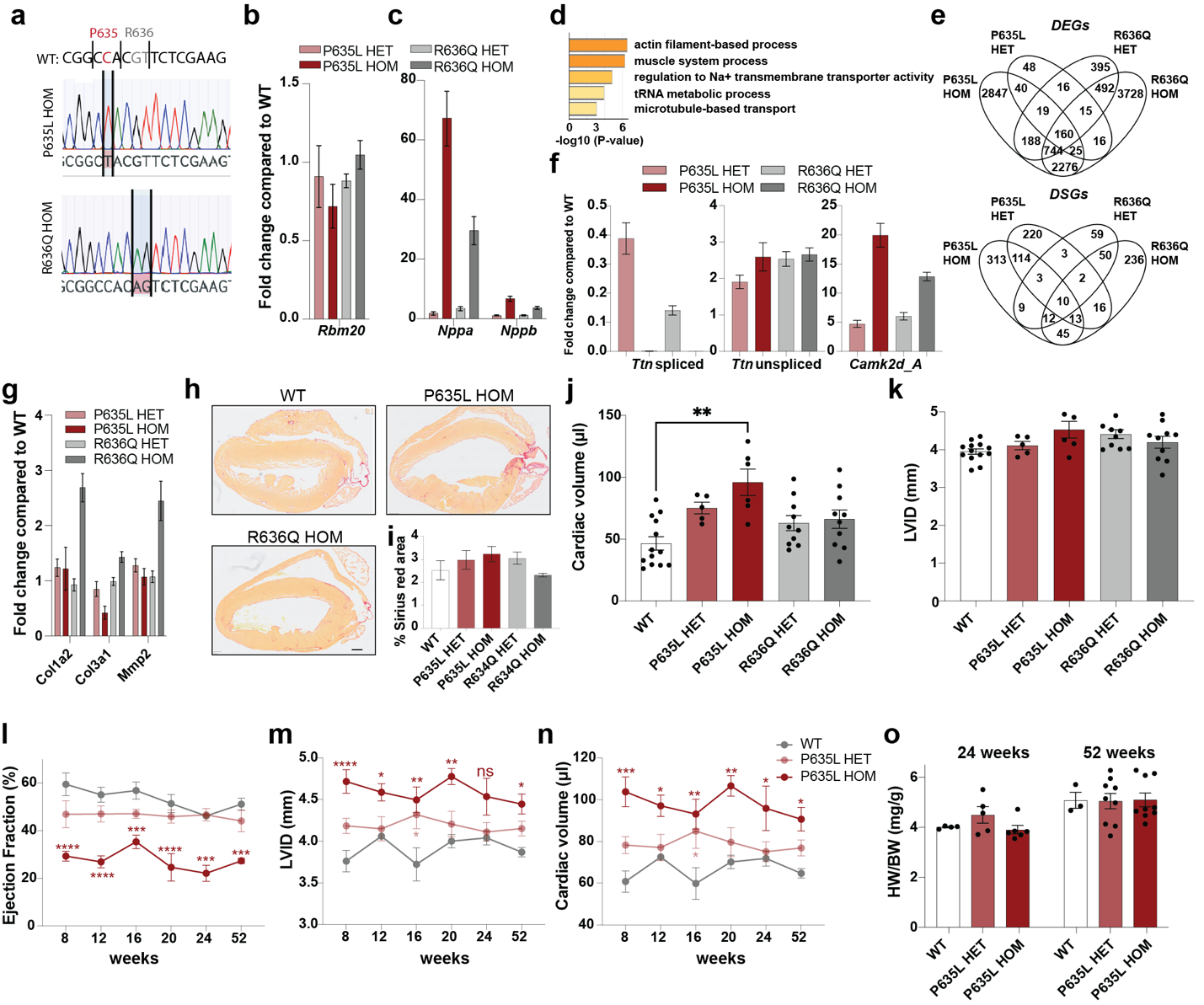

**Supplementary Figure 1: Extended characterization of P635L and R636Q mouse lines. a)** Sanger sequencing traces of first generation homozygous mutant mice used for subsequent mating and experiments. Red: C>T mutation leading to the P635L amino acid change; grey: GT>AG mutation inducing the R636Q substitution. Note that for subsequent base editing in R636Q, the CAG codon is converted to CGG which is synonymous to the WT CGT codon. **b, c)** Expression fold change compared to WT of *Rbm20* (**b**), and *Nppa* and *Nppb* (**c**). N = 5-6 mice per genotype. **d)** GO analysis (biological function) of DSGs that overlap for both P635L and R636Q HOM mice with a cut-off of p_adjust_ <0.01 and ΔPSI > 0.1. **e)** Venn diagram of significantly DEGs (up, p_adjust_ <0.05) and DSGs (down, p_adjust_ <0.01 and ΔPSI > 0.1). **f, g)** Expression fold change compared to WT of spliced and unspliced *Ttn* isoforms and *Camk2d* isoform A (**f**) or fibrosis marker genes (**g**) determined by qPCR. N = 5-6 mice per genotype. **h, i)** Representative heart tissue sections stained with Sirius Red (**h**) and quantification of Sirius Red positive area (**i**). N = 3 mice per genotype. Scale bar: 500 μm. **j, k)** Cardiac volume (**j**) and LVID (**k**) determined by narcosis echocardiography of mutant mice. N = 5-13 mice per genotype. Only significant differences are labeled. P-values obtained from one-way ANOVA with Tukey`s multiple comparison test: ** p< 0.01. Mice were 24-weeks old. **l-o)** Percentage of ejection fraction (**l**), LVID (**m**) and cardiac volume (**n**) determined by time-course narcosis echocardiography, and heart-to-body weight ratio after 24 and 52 weeks (separate cohort) (**o**) of P635L HET and HOM mutant mice. N = 5-6 mice per genotype. Asterisk indicates statistical significance compared to WT obtained by two-way ANOVA with Tukey`s multiple comparison test: **** p< 0.0001, *** p< 0.001, ** p< 0.01, * p< 0.05. Differences in P635L HET versus WT were only significant when indicated. No significant changes were found in (**i**), (**k**) and (**o**). All data was obtained in 16-week-old mice if not indicated otherwise. Error bars depict the SEM.

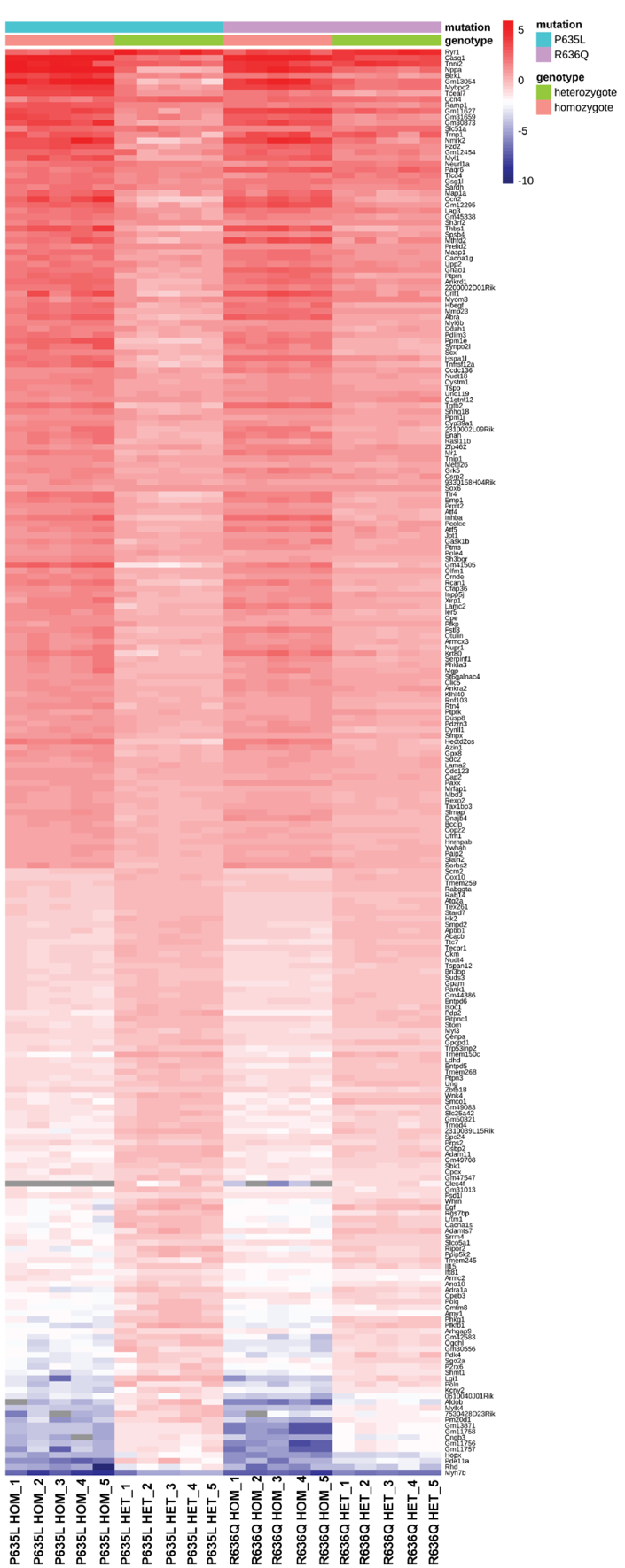

**Supplementary Figure 2: Heatmap of DEGs of P635L and R636Q mice derived from RNA-seq.** Grey bars indicate that the gene was not detected in RNA-seq data. N = 5 mice per genotype. All DEGs that overlap in P635L and R636Q HOM mice are shown with a stringent cut-off of p_adjust_ <1e^-10^.

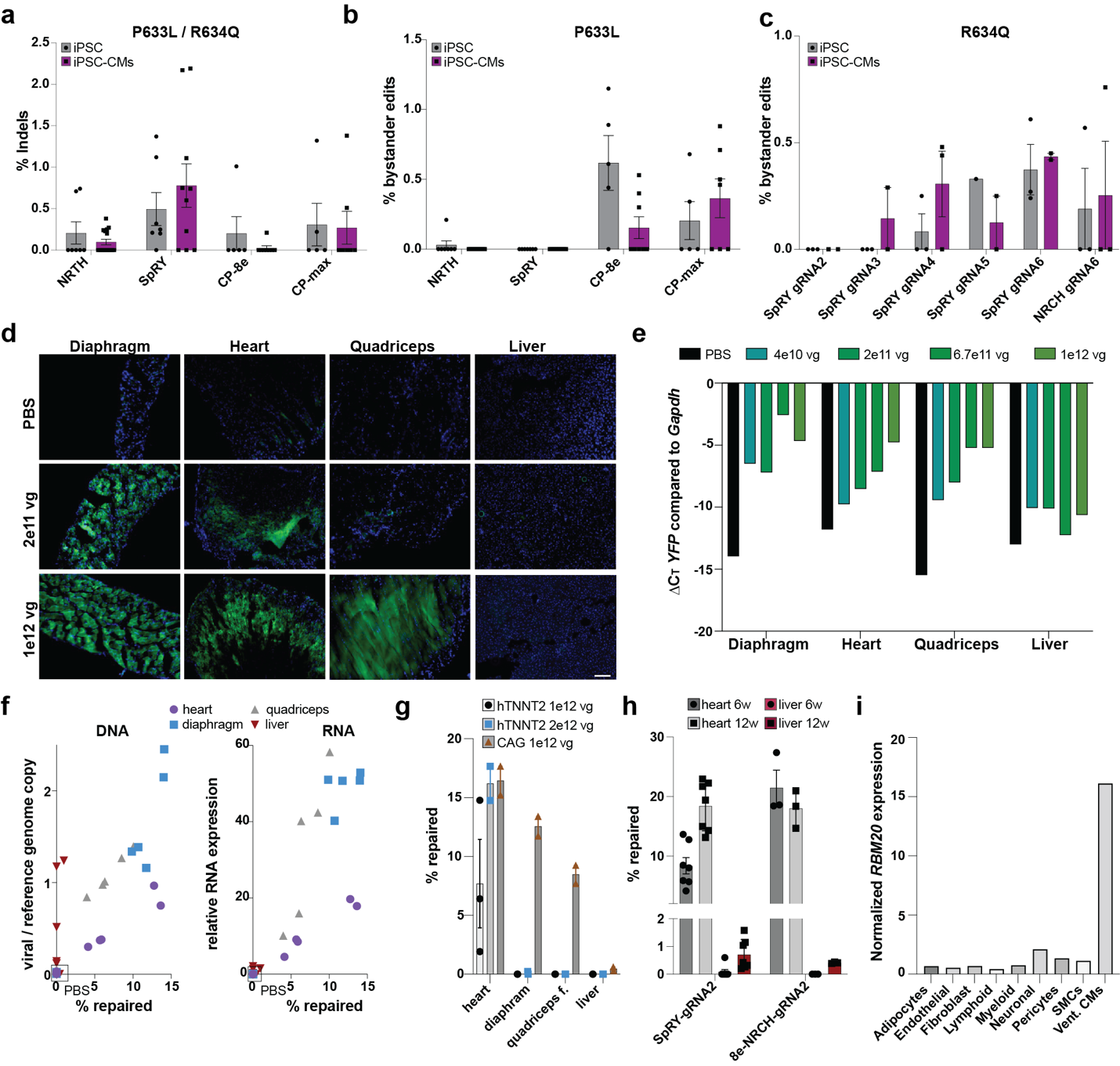

**Supplementary Figure 3: Analysis of base editing in iPSC-CMs and in mice. a-c)** Percentage of Indels (**a**) and bystander edits in P633L (**b**) and R634Q (**c**) iPSCs and iPSC-CMs. Indel formation is the summed frequency of insertions or deletions in a window of 10 bp upstream to 10 bp downstream of the gRNA binding site. Indel formation is shown combined for both mutations and gRNAs and separated by different base editors only. Bystander edits are sequences that contain extra A>G conversions within the gRNA window. **d, e)** Representative tissue sections (**d**) and RNA expression data (**e**) measuring the fluorescent transgene *YFP* delivered by AAVMYO and injected in different concentrations in WT mice. Scale bar: 100 μm. One mouse per concentration was injected. **f)** Percentage of repaired reads relative to viral copy number per diploid genome (**left**) or relative to RNA expression (**right**) determined by ddPCR in the muscle tissues heart, diaphragm and quadriceps femoris (quadriceps f.), as well as the liver. For the left panel, DNA was used as input with primers for the CMV promoter, for the right panel, RNA reverse transcribed to cDNA was used with primers for the transcribed WPRE element common in all base editor constructs. Only the SpRY-gRNA2 combination was analyzed. **g)** Editing efficacy of NRCH / gRNA2 driven by the *hTNNT2* or the *CAG* promoter. Concentration shown is the combined amount of N-and C-terminal base editor-containing AAV. **h)** Editing efficacy of SpRY and 8e-NRCH in heart and liver 6 and 12 weeks after injection in 4-week-old mice. **i)** Normalized RNA expression of RBM20 derived from single-nucleus RNA-seq of the human heart. SMCs = smooth muscle cells, Vent. CMs = ventricular cardiomyocytes. Error bars depict the SEM. Only P635L HOM were treated.

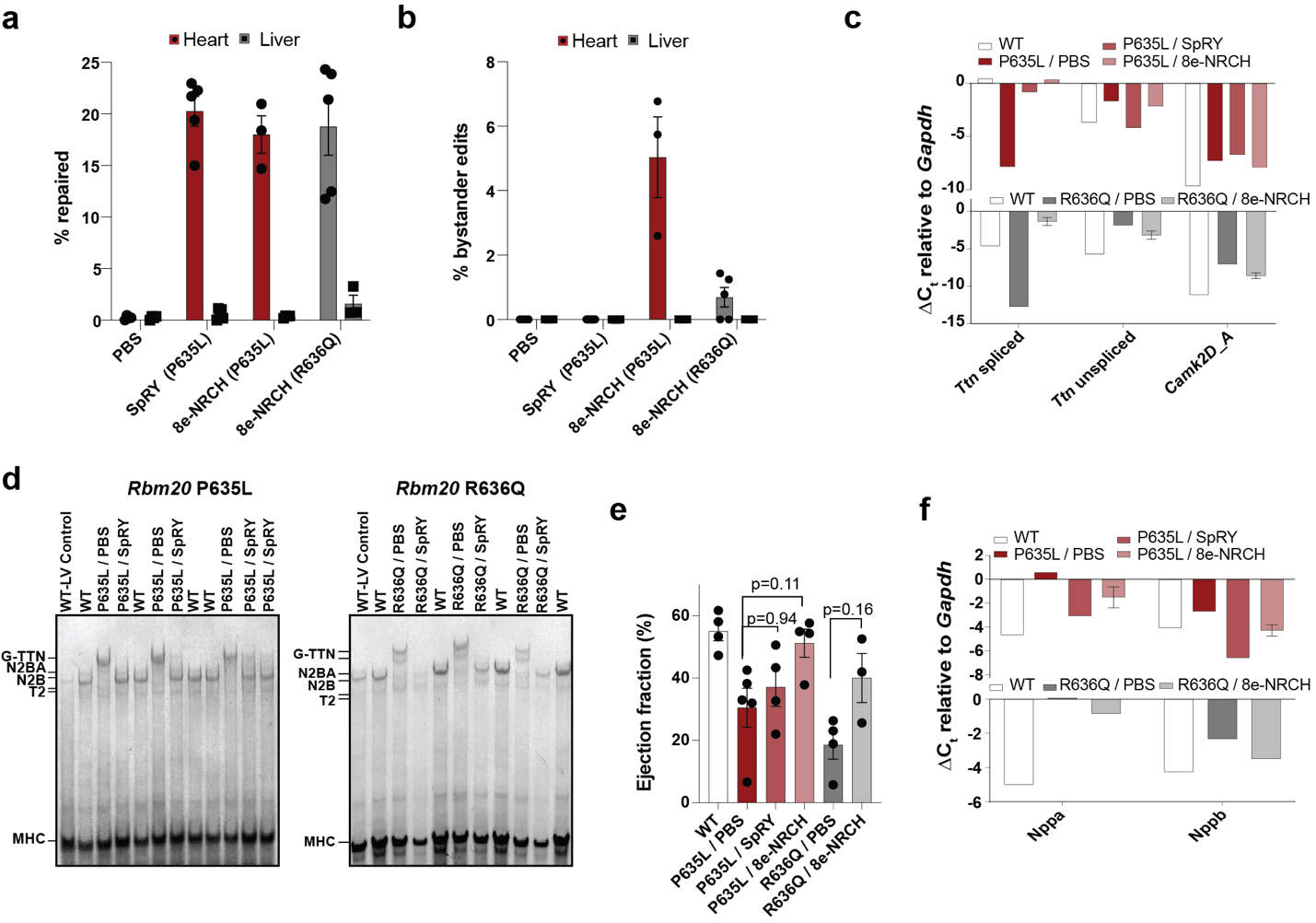

**Supplementary Figure 4: Extended phenotypic characterization of mice after AAVMYO-ABE treatment. a, b)** Allele frequency of repaired DNA (**a**) and bystander edits (**b**) in mice treated with AAVMYO-ABE. **c)** Expression of spliced and unspliced *Ttn* isoforms and *Camk2d* isoform A in WT and mutant mice treated with PBS or AAVMYO-ABE. N = 3-4 mice per condition. **d)** Vertical TTN agarose gels showing the G-N2BA, N2BA N2B, and T2 protein isoforms of TTN, and MHC as a loading control**. e)** Percentage of ejection fraction in WT or mutant mice 8 weeks after injection with PBS or AAVMYO-ABE. P-values obtained by one-way ANOVA with Tukey`s multiple comparison test shown for AAVMYO versus PBS. **f)** Expression of heart failure biomarkers *Nppa* and *Nppb* in WT and mutant mice treated with PBS or AAVMYO-ABE. N = 3-4 mice per condition. Only P635L or R636Q HOM mice were treated. Error bars depict the SEM. All data except in (**e**) were obtained 12 weeks after AAVMYO-ABE injection.

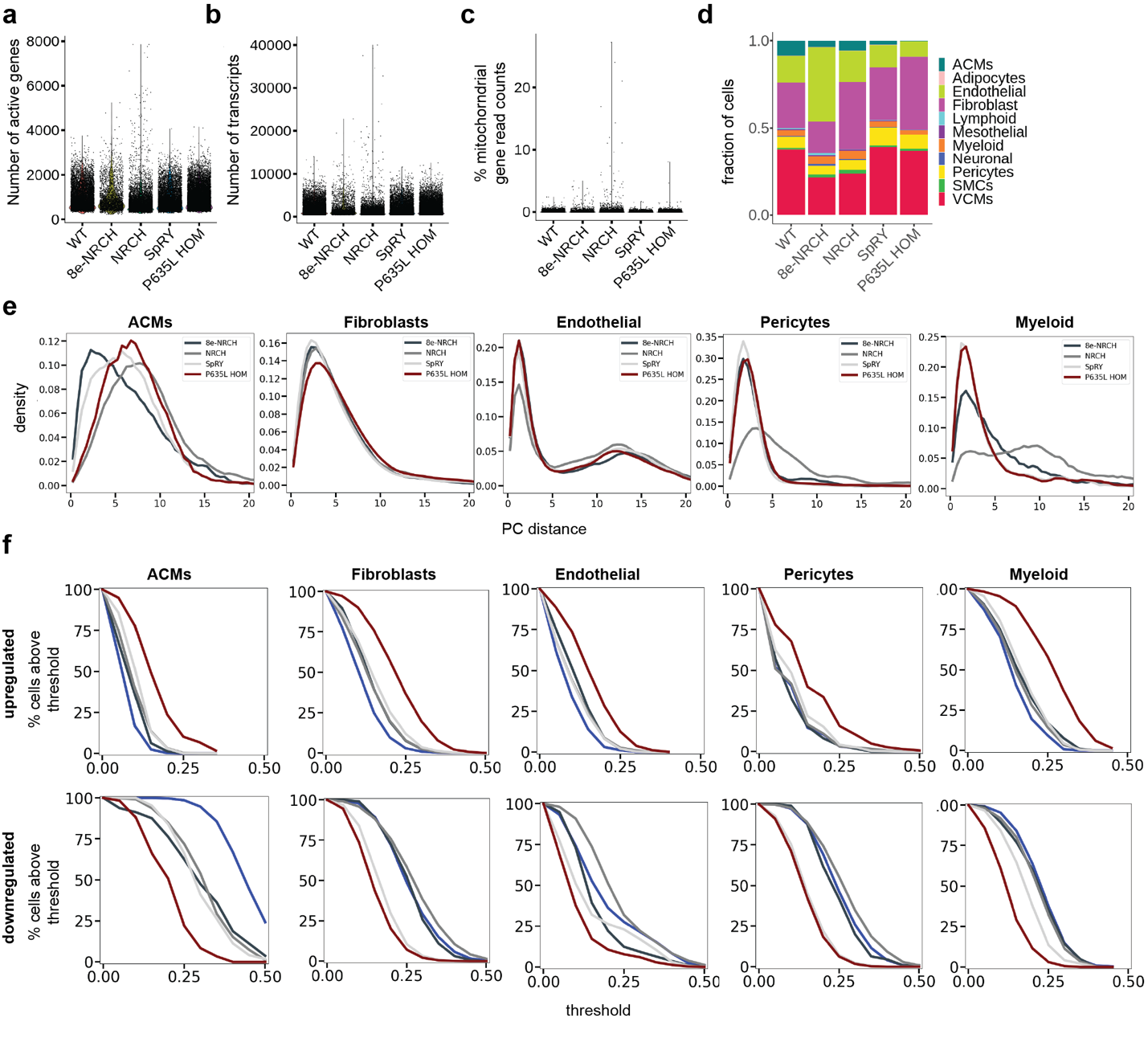

**Supplementary Figure 5: Extended snRNA-seq analysis. a-c)** Number of active genes (**a**), total transcript counts (**b**) and percentage of mitochondrial gene counts per cell (**c**) for nuclei from each condition. Two independent snRNA-seq experiments were performed for each condition except for NRCH where only one experiment was sequenced. **d)** Relative cell type distribution in WT, P635L HOM and base edited mice. **e)** Histograms depicting the distribution of pairwise Euclidean distances of depicted cell types from P635L HOM and base edited mice relative to WT upon mapping using two principal components (PC). **f)** Threshold of activity score of depicted cell types (see methods for calculation) relative to percentage of cells above the threshold for genes upregulated (upper panel) or downregulated (lower panel) in P635L HOM relative to WT.

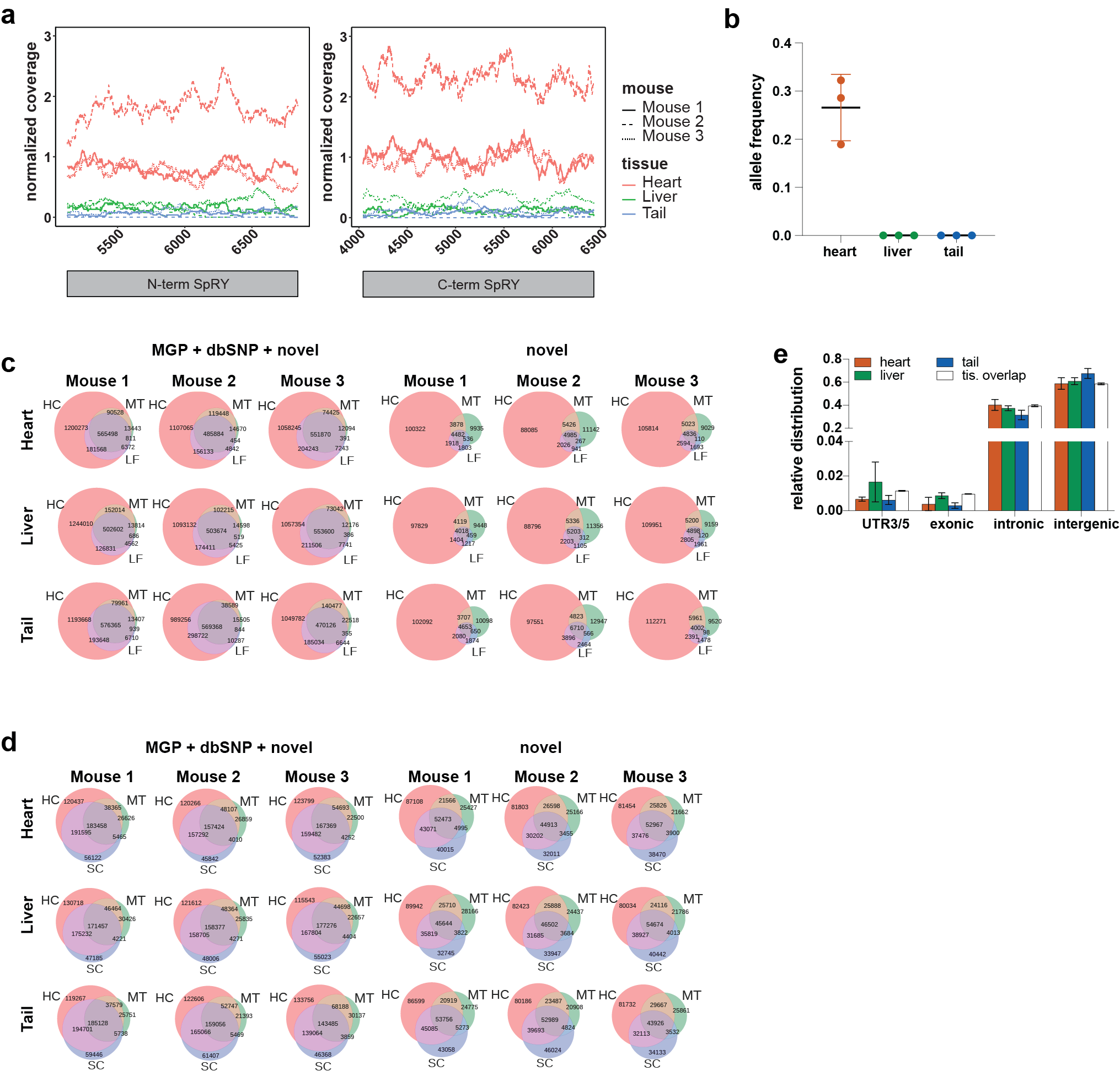

**Supplementary Figure 6:** **AAV coverage and editing events detected by WGS. a)** Normalized read coverage across autosomes of C- and N-terminal base editor sequence delivered by AAVMYO. **b)** Allele frequency of the P635L A>G nucleotide conversion. **c, d)** All (left) and novel (right) SNVs (**c**) or Indels (**d**) called by four variant callers (HC: HaplotypeCaller, MT: Mutect2, LF: Lofreq, SC: Scalpel). **e)** Mean relative distribution of tissue-specific and common variants within coding or non-coding regions of the genome. Tissue overlap represent variants that were common to all three tissues.
